# Protists as mediators of complex microbial and viral associations

**DOI:** 10.1101/2024.12.29.630703

**Authors:** Frederik Schulz, Ying Yan, Agnes K.M. Weiner, Ragib Ahsan, Laura A. Katz, Tanja Woyke

## Abstract

Microbial eukaryotes (aka protists) are known for their important roles in nutrient cycling across different ecosystems. However, the composition and function of protist-associated microbiomes remains largely elusive. Here, we employ cultivation-independent single-cell isolation and genome-resolved metagenomics to provide detailed insights into underexplored microbiomes and viromes of over 100 currently uncultivable ciliates and amoebae isolated from diverse environments. Our findings reveal unique microbiome compositions and hint at an intricate network of complex interactions and associations with bacterial symbionts and viruses. We observed stark differences between ciliates and amoebae in terms of microbiome and virome compositions, highlighting the specificity of protist-microbe interactions. Over 115 of the recovered microbial genomes were affiliated with known endosymbionts of eukaryotes, including diverse members of the Holosporales, Rickettsiales, Legionellales, Chlamydiae, Dependentiae, and more than 250 were affiliated with possible host-associated bacteria of the phylum Patescibacteria. We also identified more than 80 giant viruses belonging to diverse viral lineages, of which some were actively expressing genes in single cell transcriptomes, suggesting a possible association with the sampled protists. We also revealed a wide range of other viruses that were predicted to infect eukaryotes or host-associated bacteria. Our results provide further evidence that protists serve as mediators of complex microbial and viral associations, playing a critical role in ecological networks. The frequent co-occurrence of giant viruses and diverse microbial symbionts in our samples suggests multipartite associations, particularly among amoebae. Our study provides a preliminary assessment of the microbial diversity associated with lesser-known protist lineages and paves the way for a deeper understanding of protist ecology and their roles in environmental and human health.

## Introduction

Protists, microeukaryotes that are not fungi, plants or animals^1^, play instrumental roles in global ecosystems where they contribute to nutrient cycling and shape structure and function of microbial communities^2,3^. Heterotrophic protists are well known for their role in grazing, a process in which other microbes are being taken up through phagocytosis and digested^4^. This turnover of microbial biomass ultimately makes nutrients available to higher trophic levels^2^. Grazing isn’t the sole process that protists are involved in. Both heterotrophs and autotrophs (i.e. unicellular algae) play central roles in biomineralization^5^, while autotrophs and mixotrophs contribute to organic carbon fixation^6,7^.

Recent studies suggest that protists may harbor complex microbiomes^8^. For example, some amoebae have evolved strategies to maintain a regime of bacteria as part of their microbiome as food to ensure a constant nutrient supply^9^. Some of the protist-associated bacteria are resistant to digestion^10^ and may even be able to replicate inside their eukaryotic host cells^11^.

This ability might lead to host dependency^12^ or even protection against pathogens^1^^3^. Further, amoebae represent natural reservoirs for a wider range of human pathogens^14^. This complexity could have far-reaching implications, not only for the surrounding microbial communities and nutrient cycling but also for ecosystem, animal, and plant health.

Despite their importance, our insights into the roles of protists in ecosystems and particularly their interactions with associated microbes and viruses remain limited to a few well-studied groups such as *Acanthamoeba* and *Paramecium^15–17^*. These have received attention due to their medical relevance and their ability to be cultivated under axenic or monoxenic conditions. This leaves broad gaps in our understanding of diversity, function and associations of lesser studied protist lineages even though they make up most branches in the eukaryotic tree of life^11,18–20^.

To address these limitations, we collected over 100 individual cells of diverse microbial eukaryotes directly from the environment, including ciliates and testate (i.e. shell-building) amoebae. Most of these organisms are understudied and have not been successfully maintained in culture. Using cultivation-independent single cell isolation, whole genome amplification and genome resolved metagenomics, as well as single cell transcriptomics, we provide insights into the unique composition of the protist microbiome and virome. We uncover associations with a wide range of putative pathogens, including both microbial and eukaryotic symbionts and viruses that infect protists and their associated microbes. Our findings underscore the role of protists as mediators of complex microbial and viral infections in the environment and shed light on the intricate roles that these organisms play within ecological networks.

## Results & Discussion

### Protist microbiome composition and diversity

The metagenomic binning of sequences from 104 single amplified genomes (SAGs) belonging to three testate amoeba and eight ciliate species (Figure 1a, Supplementary table 1) yielded a total of 724 prokaryotic metagenome-assembled genomes (MAGs; Supplementary tables 1,2).

**Figure 1.**
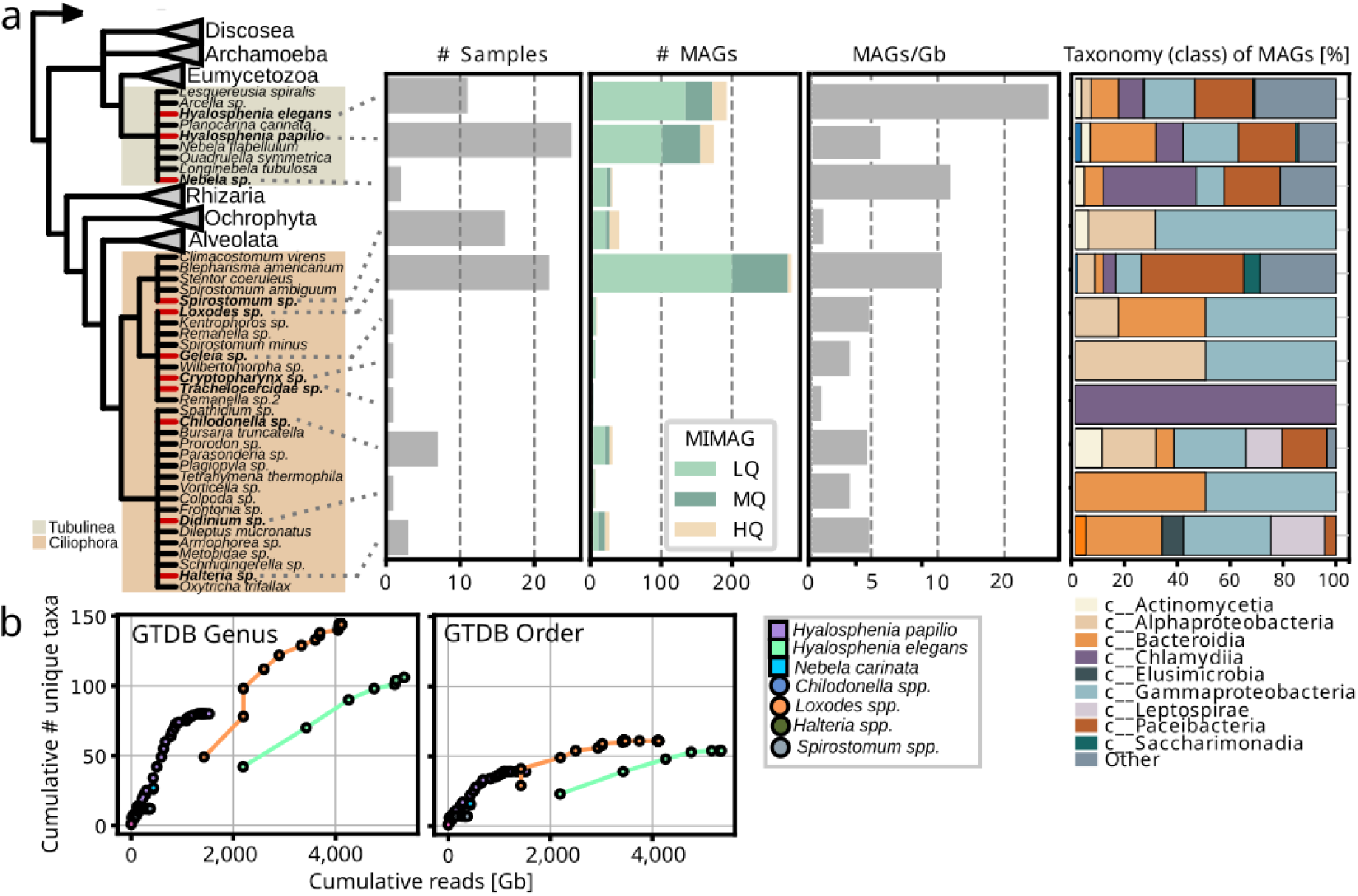
Amoeba (Tubulinea) and ciliate (Ciliophora) lineages sampled in this study. (a) Simplified phylogenetic tree indicates protist lineages sampled in this study (in bold with red branches), bars indicate number of samples sequenced, total number of MAGs per protist lineage and number of MAGs per Gb sequenced. The right panel shows taxonomic distribution of bacterial and archaeal MAGs associated with sampled protists. (b) Collector’s curves comparing the overall richness of bacterial and archaeal MAGs at different taxonomic levels for each protist lineage.

According to MIMAG standards^21^, 442 were of low, 209 of medium and 76 of high quality. Sequencing depth varied between samples but the greatest number of MAGs per gigabase (Gb) of reads was recovered from the amoeba *Hyalosphenia elegans* and the ciliate *Loxodes* sp., the latter of which was deeply sequenced (Figure 1a). *Hyalosphenia elegans* showed a much higher recovery rate of MAGs per Gb of sequence data compared to its sister species, *Hyalosphenia papilio*. This trend was also visible in the overall microbiome diversity; *Hyalosphenia elegans* was associated with a greater number of detected bacterial and archaeal phyla (n=16), orders (n=52) and genera (n=52), whereas despite higher sequencing depth *Hyalosphenia papilio* plateaued at 15 phyla, 35 orders and 80 genera (Figure 1b). This may reflect the relative sizes of these organisms as more of the genomic material may be host for the larger *H. papilio* (100-150µm) compared to *H. elegans* (80-120 µm) and also the presence of additional microalgal symbionts in *H. papilio*^22^. Overall, microbiome diversity varied strongly among the sampled lineages; for example, MAGs recovered from *Spirostomum* sp. belonged to just two different bacterial phyla, while those from *Loxodes* sp., isolated from the same low pH environments as the testate amoebae, belonged to up to 16 different bacterial phyla, which corresponded to at least 60 orders and 145 genera (Figure 1). When comparing the protist microbiomes, it became apparent that ciliates and amoebae have distinct microbiome compositions (Figure 1a). Distribution of bacterial classes was more similar between amoebal lineages and more dissimilar between ciliates, with the freshwater *Loxodes* and the marine Trachelocercidae having the least proportion of shared taxa with other ciliates.

### Protist microbiomes harbor microbes across known endosymbiont clades

A large proportion of recovered taxa belong to groups of known host-associated bacteria with a facultative or even obligate intracellular lifestyle. Specifically, 115 prokaryotic MAGs grouped with known bacterial endosymbionts and 258 with putative symbionts (Patescibacteria)(Figure 2). Alphaproteobacterial endosymbionts were exclusively found in association with ciliates, particularly Megaira and Caedimonadales in *Spirostomum*, and Paracaedibacterales with *Loxodes*, *Chilodonella* and *Halteria* (Figure 2). Additionally, several novel and currently uncharacterized lineages within the order Rickettsiales were detected within *Loxodes* and, to a lesser extent, *Chilodonella*. Three of the sampled Loxodes cells contained bacteria that grouped in the gammaproteobacterial family UBA6186 together with Azoamicus ciliatocola, a bacterial endosymbiont of ciliates with cosmopolitan distribution^23^ that has been shown to generate energy for its host by denitrification^24^. Members of Holosporales and Megaira have previously been associated with different ciliates^25^, but none of the other lineages have been identified as ciliate endosymbionts thus far. In both species of *Hyalosphenia* sampled in this study, likely host-associated gammaproteobacteria of the family *Francisellaceae* were found. Further, Diplorickettsia were present in *Hyalosphenia elegans*, along with Coxiellales-related bacteria in the ciliate *Cryptopharynx*, and members of Legionellales were present in the ciliates *Didinium* and *Loxodes* as well as the testate amoeba *Hyalosphenia papilio*. For *Didinium* and *Cryptopharynx*, gammaproteobacteria were the only associated putative symbionts. Another group of protist symbionts, the phylum Dependentiae, had members from four different families associated with amoebae and ciliates sampled in this study; *Chromulinavoraceae* were exclusively found with *Hyalosphenia papilio (an amoeba that harbors green algal symbionts)*, while other families were mixed between *Hyalosphenia* species and *Loxodes*.

**Figure 2.**
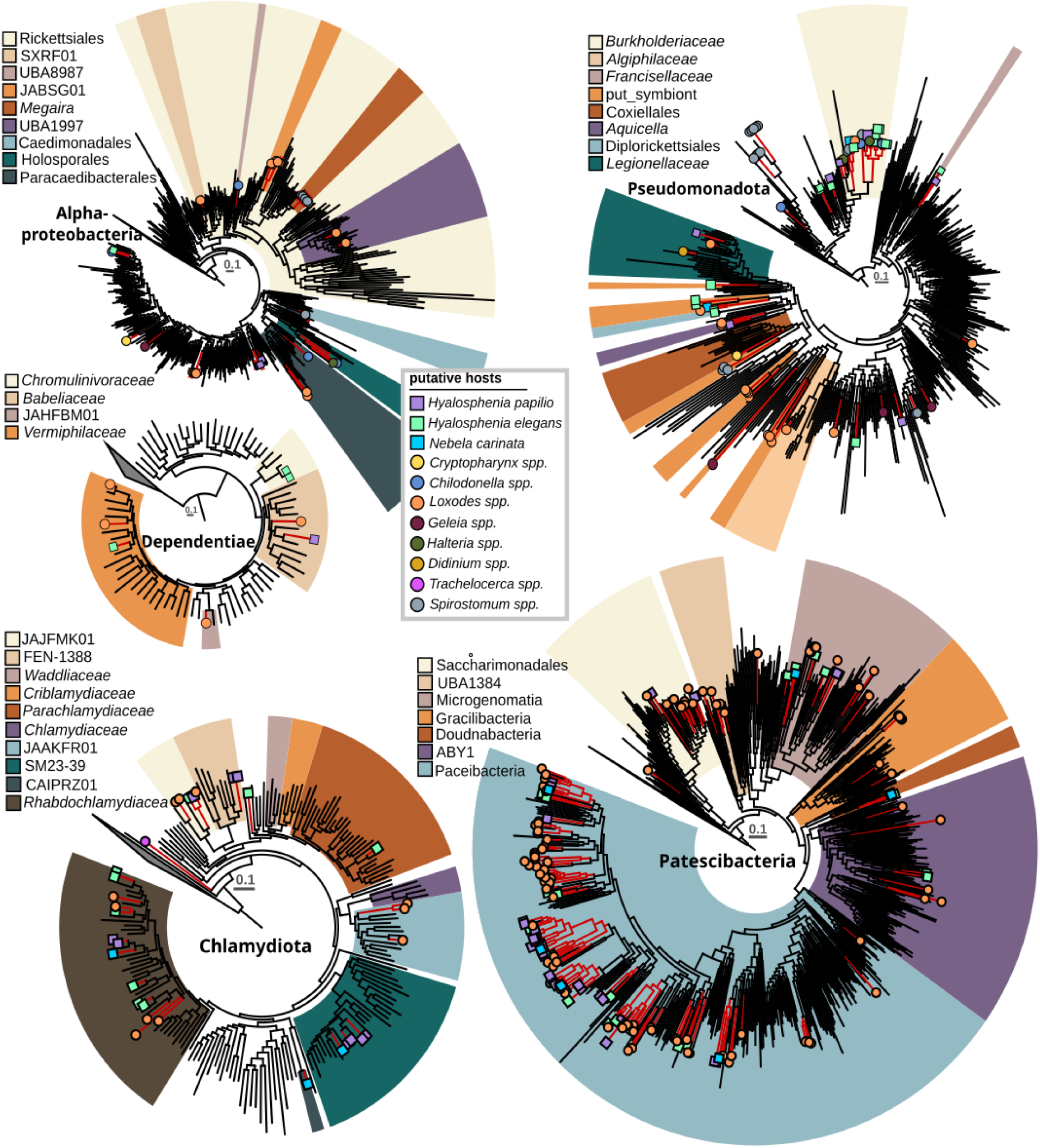
Amoeba and ciliate associated bacteria affiliated with known and suggested bacterial endosymbiont clades. Phylogenetic trees indicate the position of putative symbiont MAGs (red branches) that were recovered from protist microbiomes generated in this study. Colored shapes at the terminal branches show the protist lineages the MAGs were associated with. Each tree represents a phylum (Chlamydiota, Dependentiae, Patescibacteria) or class (Alphaproteobacteria, Pseudomonadota) and subclades that were previously shown (in Chlamydiota, Dependentiae, Alphaproteobacteria, Pseudomonadota) or suggested (in Patescibacteria) to be host-associated are highlighted with colored wedges.

This is the first time Dependentiae have been identified as potential ciliate symbionts. One of the best studied symbiont clades is the phylum Chlamydiota, known to infect a wide diversity of eukaryotic hosts^17^. Here, we identified 56 chlamydial MAGs associated with diverse ciliates and amoebae. Four family-level lineages that consist solely of metagenome-assembled genomes were associated with either *Hyalosphenia* (f FEN-1388), *Loxodes* (f JAAKFR01), *Hyalosphenia* and *Loxodes* (f JAJFMA01), or *Hyalosphenia* and *Nebela* (f SM23-39). Further, the only bacterial symbiont which was found associated with *Trachelocerca* was a highly divergent member of the Chlamydiota, potentially representing a novel family or even order-level lineage without any closely related relatives (Figure 2). The amoeba *Hyalosphenia* was found to be associated with Parachlamydiacaea, a group previously shown to infect different amoebae, particularly *Acanthamoeba castellanii*. In *Acanthamoeba* it confers protection against giant virus infection^26^ but it is also associated with disease in humans and other animals^27^. Most chlamydial MAGs recovered in this study were associated with four different ciliates (*Loxodes*) and amoebae (*Hyalosphenia* and *Nebela*). These were affiliated with Rhabdochlamydiacae, a group previously shown to be predominantly associated with insects and other metazoans^28^.

### Diverse Patescibacteria make up a large fraction of the protist microbiome

In addition to members of well-known intracellular bacteria, we recovered 258 MAGs (25 high quality, 79 medium quality and 154 low quality) representing members from all major groups of the phylum Patescibacteria, mainly associated with *Loxodes*, *Nebela*, and *Hyalosphenia papilio* and, to a lesser extent, with *Hyalosphenia elegans*, *Halteria*, and *Chilodonella* (Figure 2). Given the reduced genomes of Patescibacteria and other features that may underlie host interaction, it’s plausible that some or all of these might be closely associated with amoebae and ciliates. This aligns with a previous study that provided experimental evidence of an uncharacterized Parcubacterium as an intracellular bacterium in the ciliate *Paramecium* sp.^29^. However, reports also exist of association with other bacteria^30,31^ and of a potential free-living lifestyle for Patescibacteria^32,33^. Our results indicate some patterns of co-occurrence, such as clades of Paceibacteria composed of MAGs derived from *Loxodes* (Figure 2). However, the high overall diversity of Patescibacteria MAGs and absence of clear host-specificity pattern, hampered any predictions in regards to endosymbiosis.

### Giant viruses and virophages are frequently found in protist microbiomes and genes transcribed *in situ*

The microbiomes of the ciliates and amoebae sampled in our study did not only contain sequences of various host-associated bacteria but 82 giant viruses metagenome assembled genomes (GVMAGs) (Supplementary table 3). Taxonomic identification with gvclass^34^ and phylogenomic analysis revealed that these GVMAGs belonged to diverse lineages within the viral phylum Nucleocytoviricota^16^ (Figure 3a). Specifically, those associated with *Hyalosphenia* belonged to several orders, including Asfuvirales, Pandoravirales, Algavirales, and Imitervirales. Viruses associated with *Loxodes* were highly diverse; however, those linked to *Hyalosphenia papilio* and *Chilodonella* were confined to a few clades within Imitervirales. In previous studies giant viruses have not been found to directly infect ciliates^35^. However, the frequent presence of diverse giant viruses in the ciliates *Loxodes* and *Chilodonella* sampled here suggests members of Ciliophora as underappreciated potential hosts for these viruses. For samples where sequences from more than one eukaryote were found, inferring sequence-based putative associations is challenging. For example, members of Algavirales might more likely infect the green algae^36^ that are symbiotic to *Hyalosphenia* and were also detected in the same samples, rather than the *Hyalosphenia* itself. Further, it has been shown that giant viruses are frequently ingested as food^37^. Such uptake may not lead to an infection in amoebae and ciliates and viruses may accumulate in the cytoplasm, or in some cases multiple highly similar viruses are taken up at the same time^38,39^.

**Figure 3.**
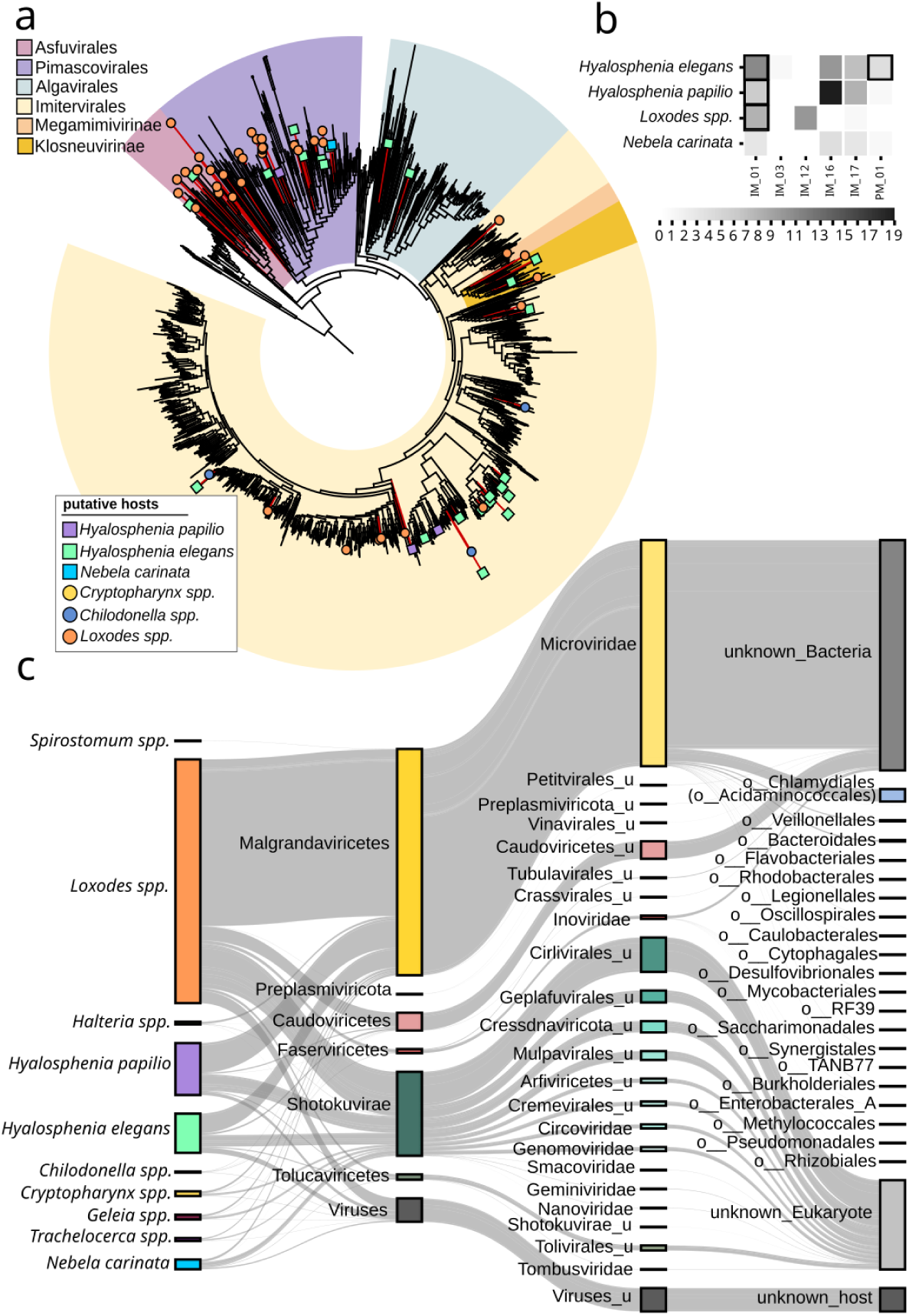
Viral sequences found in ciliate and amoeba microbiomes. (a) Phylogenetic tree of the Nucleocytoviricota. Red branches indicate giant virus genomes recovered in this study. Color of circles (ciliates) and squares (amoeba) at terminal branches correspond to protist lineages. Colored wedges highlight order level groups in the Nucleocytoviricota. (b) Nucleocytoviricota families for which genes were found to be expressed, as based on single cell transcriptomes of four protist lineages. Numbers indicate the number of protist single cells associated with members of the same active Nucleocytoviricota family. Fields with a black outline indicate pairs which were also identified in the genome data. In addition to these viral transcripts there were others that could be assigned viruses of the families IM_03, IM_12, IM_16 and IM17, but none of these were found in the SAG data (c) Sankey diagram linking non-Nucleocytoviricota viral contigs of high completeness to protists and predicted hosts. Segments on the left are colored based on the protist lineages, segments in the two center columns represent viral lineages on the order and family level, segments on the right correspond to predicted hosts.

To better understand if some of the detected giant virus lineages are actively infecting the protists, we analyzed single cell transcriptomics on similar various amoebae and ciliates directly isolated from our sampling sites (Supplementary Table 1). Using these data, we were able to confirm gene expression for viruses of the Imiterviales family IM_01 (Mesomimiviridae) in several *Hyalosphenia elegans*, *Hyalosphenia papilio* and *Loxodes* cells. Further, we found genes of members of the Pimascovirales family PM_01 expressed in *Hyalosphenia elegans*. In-depth experimental assessment of the protist lineages sampled here, which however are challenging to maintain in the lab, will be required to fully establish a direct connection between giant viruses and a particular protist host.

In addition to giant viruses, we were also able to recover sequences of 33 virophages from the sorted protists (Supplementary Table 4), of which 25 had sufficient number of virophage hallmark genes to be placed into the Lavidaviridae taxonomic framework (described in ^40^). 27 out of 33 virophage sequences were found in protist microbiomes which also contained giant virus genomes. Virophages are known to integrate into their host genomes and become activated when the host encounters giant viruses, which they parasitize, potentially offering protection against giant virus infection^41^. All virophages recovered here were on contigs with a length of 5-26kb which is the typical genome size range of known virophages^42,43^ with none found on longer contigs or surrounded by protist genes. Virophage genes were not found actively expressed in metatranscriptome data from independently sorted similar protists. Nevertheless, our findings suggest that the virophages are probably not integrated into the protist genome, but rather actively engaging in virus-virus interactions.

### Protists are hot spots of DNA virus diversity

All sampled protists in our study were associated with a large number of other viruses (Figure 3c; Supplementary table 5). Most of these belonged to the subfamily Gokushovirinae from the family Microviridae in the order Malgrandaviricetes, which are ssDNA viruses that typically have small genomes and are known to infect host-associated bacteria^44^. Host prediction based on iPHoP^45^ indicated a broad range of potential hosts, including Legionellales, Coxiellales, Burkholderiales, Acidaminococcales, and others (Figure 3c). To a lesser extent, we found members of the order Caudoviricetes, which are diverse tailed dsDNA viruses associated with free-living bacteria. We also identified other viruses that we could not taxonomically classify but that were predicted to infect intracellular bacteria in the order Chlamydiales (Figure 3c). Additionally, we found numerous ssDNA viruses from the Shotokuvirae, most of which belonged to the Cressdnaviricota orders Arfiviricetes and Repensiviricetes, all known to infect a wide range of eukaryotic hosts^46^. Host prediction for eukaryotic viruses is less advanced than for bacterial viruses, so it’s not entirely clear which of the detected viruses may infect the protists or associated eukaryotes, or whether these viruses adhere to the protist surface or reside in the cytoplasm or food vacuoles prior to being degraded. Given the diversity and abundance of detected viruses, it’s conceivable that some indeed infect protist hosts.

### Complex multipartite associations in protists microbiomes

Our single cell study suggests that multipartite associations among protists, bacterial symbionts, giant viruses, and other viruses are prevalent (Figure 4). This is particularly true for amoebae SAGs for which sequences from giant viruses, Chlamydia, Dependentiae and Gammaproteobacteria affiliated with known intracellular bacteria frequently co-occurred. In ciliate SAGs, the co-occurrence pattern differed and multipartite associations were mainly predicted in *Loxodes* sp. consisting of giant viruses and chlamydial and alphaproteobacterial symbionts, and to a lesser extent, Dependentiae or Gammaproteobacteria. There was only a single case (*Chilodonella* sp.) of a predicted multipartite association that involved three or more interaction partners. For other ciliate lineages we did not identify multiple interaction partners (Figure 4). The overall lower complexity of sequences from microbial symbionts and giant viruses in different ciliate samples can potentially be attributed to two factors: first, different grazing preferences compared to amoebae^47^ and second, the absence of other associated microeukaryotes in the same samples. In contrast, most amoebae were associated with sequences from smaller, often flagellated protists, such as kinetoplastids and chrysophytes. In the case of testate amoeba, washing of their shells is more difficult to achieve compared to ciliates or naked amoebae and may have hindered the complete removal of attached smaller eukaryotes. Additionally, green algae were frequently detected, especially in *Hyalosphenia papilio* which is known to be associated with endosymbiotic *Chlorella*^22^. Notably, while Chloroviruses are known to associate with *Chlorella*, we did not recover any giant viruses from *Hyalosphenia* samples containing *Chlorella*. In contrast, we sampled three *Hyalosphenia elegans* cells that were not associated with any other eukaryotes but each associated with multiple giant viruses. Next, we tested how different factors shape the uniqueness and richness of distinct microbial and viral groups within protist microbiomes. Specifically, the sampling site was a key driver of uniqueness for free-living bacteria and giant viruses (Nucleocytoviricota), while host taxonomy most strongly influenced uniqueness in putative endosymbionts and Patescibacteria. In contrast, microbiome richness was mainly linked to host taxonomy and sequencing depth. Morphology, cell size, and lifestyle had moderate but variable effects. These findings reinforce that both host traits (e.g., taxonomy, morphology) and environmental conditions (e.g., sampling site) collectively shape the complexity of protist-associated microbial and viral communities. Taken together, our analysis supports the notion that protists serve as powerful drivers of multipartite interactions, and are tightly linked to diversity, specificity, and ecological roles of their bacterial and viral partners.

**Figure 4.**
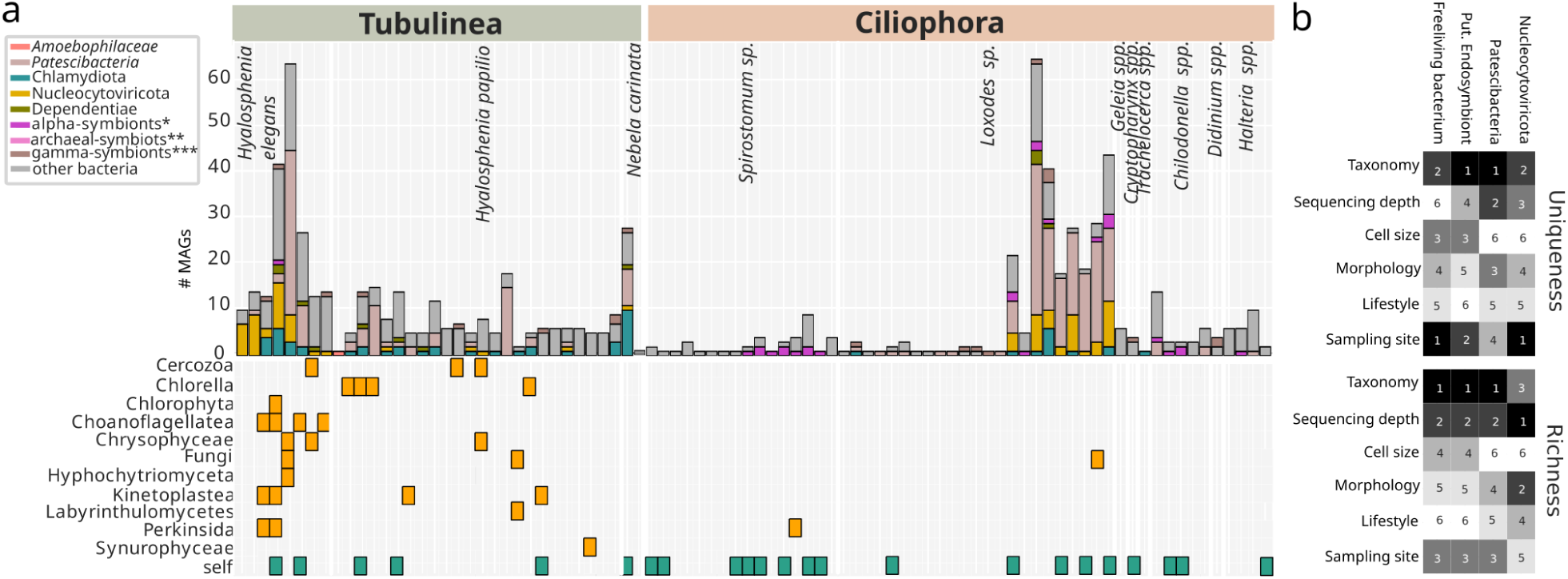
Multipartite association in ciliate and amoeba microbiomes, as based on SAG sequences. (a) The upper panel shows the distribution of different symbiont lineages, giant viruses and free living bacteria in the individual assemblies of the sampled protists. The lower panel indicates other eukaryotes that were associated with the respective data sets. *self detection of protist hosts based on 18S rRNA gene screening. * Rickettsiales, Holosporales, Paracaedibacterales, Caedimonadales; **Methanomicrobiales; ***Legionellales, Coxiellales, Diplorickettsiales, *Algiphilaceae*, *Aquicella*, *Franciscellaceae*. (b) Ranked impact of factors on uniqueness (upper heatmap) and richness (lower heatmap) of freeliving bacteria, putative endosymbionts, Patescibacteria and Nucleocytoviricota, with “1” (black) having the strongest impact and “6” (white) having the least impact on the respective factor.

## Conclusion

In previous studies, protists have been identified as hosts for diverse lineages of new endosymbionts, but such findings often relied on morphological descriptions following isolation^11,48^. Here, we used cultivation-independent sequencing approaches to sample a diversity of uncultivable protists. In our more than 100 datasets, the diversity of microbes affiliated with known microbial symbionts is unparalleled, as is the diversity of identified giant viruses and other smaller viruses, such as Arfiviricetes and Repensiviricetes that infect eukaryotes and Gokushovirinae that may infect associated symbionts. The frequent co-occurrence of sequences from diverse giant viruses, with genes found to be expressed *in situ*, and microbial symbionts likely representing multipartite associations, is particularly intriguing. Previously, *Acanthamoeba* and ciliate species have been highlighted as evolutionary melting pots and potential training grounds for pathogens of multicellular eukaryotes^49–51^. Our findings provide strong support for this hypothesis and call for further experimental work to study the microeukaryotic microbiome and virome. Understanding their roles in shaping protist populations and surrounding microbial communities, not only from an evolutionary perspective, but also in the ecological context of contributing to ecosystem dynamics through nutritional symbioses and pathogenicity, is crucial.

## Materials and Methods

### Sample collection

Individual ciliates were isolated from environmental samples obtained from a range of different sampling sites (Supplementary table 1). Amoeba were collected in low-pH bogs and fens and washed off the moss that they inhabit using prefiltered (2 µm filter) bog water after size-selecting over a 300 µm filter to discard large plant material. Arcellinida testate amoebae were then picked under an inverted microscope from the water samples using hand-held glass pipettes. They were transferred to a microscope slide with a drop of freshly filtered bog water in an attempt to wash off obvious contamination sticking to the outside of the shell. Each individual was photo documented and then transferred to a 0.2 ml tube for transcriptome/genome amplification. We sampled ciliates either from a small low pH (pH ∼4.5) pond within a local fen or from the intertidal zone of a sandy beach. Samples were filtered over an 80 µm mesh (for sandy samples) or directly poured into small Petri dishes (for pond samples). Ciliates were then observed and hand-picked with glass pipettes under an inverted microscope. Cells were washed by passing through slides of *in situ* water 2-3 times on depression slides to remove obvious surrounding contaminants (e.g. other non-target micro-eukaryotes, sediment particles). Each individual was diluted with nuclease-free water or prefiltered (0.2 µm filter) *in situ* water preceding single cell transcriptome/genome amplification.

### Whole genome and transcriptome amplification and sequencing

Whole genome amplifications of individual cells were performed using the Repli-g Single-Cell Kit (Qiagen, cat. 150345) following the manufacturer’s instructions. Most samples we incubated for the recommended 8 hours, whereas for a few small ciliates with low expected DNA content (e.g. *Cryptopharynx* spp. and *Wilbertomorpha* spp.), we extended the incubation time to 10 to 12 hours. Single-cell whole transcriptome amplifications were carried out using the SMART-Seq v4 Ultra Low Input RNA Kit for Sequencing (Clontech, cat. 634895, 634896) according to the manufacturer’s instructions. Products of both, genome and transcriptome amplifications, were purified using the Ampure XP for PCR Purification system (Beckman Coulter). A Qubit 3.0 fluorometer (Invitrogen) was used to measure the DNA concentration. Sequencing libraries for the transcriptome samples were prepared using the Nextera XT DNA Library Preparation Kit (Illumina). Library preparation for the genomic samples as well as the high-throughput sequencing of all libraries (both for genomes and transcriptomes) was carried out at the genome sequencing center at the University of California, San Diego, or at the Institute for Genome Sciences at the University of Maryland, Baltimore. A HiSeq 4000 (Illumina) sequencing platform was used for the majority of samples. The only exceptions are seven genomic samples, six from *Hyalosphenia elegans* and one from *Loxodes* sp., that were sequenced on a NovaSeq 6000 sequencing system. Samples were sequenced at 10 to 80 million reads per sample (Supplementary table 1).

### Sequence data quality control and assembly

The 150 bp paired-end raw sequencing reads were quality checked using FastQC^52^ and the BBMap toolkit^53^ was used to remove adapters and trim poor-quality reads. Given that our single cell ‘omics data from uncultivable protists tend to be complex and lack reference genomes/transcriptomes, we chose to apply more stringent trimming parameters. Transcriptome reads were trimmed with “trimq = 24, minlen = 100” and genome reads with “trimq = 28, minlen = 125”. Read normalization was performed with bbnorm prior to the assembly. De novo assemblies were then performed using SPAdes(v3.14.1)^54^) using the option -sc. Further, CrossBlock from the BBTools software package^53^ was used with default settings to minimize the effects of cross-talk between assemblies of multiplexed libraries (Supplementary table 6).

### Metagenomic binning, bin QC, gene calling, and symbiont prediction

Contigs were filtered at 2kb length and subsequently organized into genome bins based on tetranucleotide sequence composition with MetaBat2^55^ yielding 6,510 MAGs with an assembly size of above 20kb. Completeness and contamination was estimated with CheckM1^56^, CheckM2^57^ and a hmmsearch (v3.1b2) using a set of 56 universal single copy panorthologs (UNI56)^58^ and taxonomy inferred with GTDB-tk^59^. MAGs were retained if they had a CheckM1 and CheckM2 contamination of below 10% and at least 5 of UNI56 markers^58^ or were classified as Nucleocytoviricota with gvclass (https://github.com/NeLLi-team/gvclass). Genecalling was performed with prodigal^60^ using the -p meta option.

### Impact of different factors on uniqueness and richness of the protist microbiome

We first grouped MAGs by library and calculated richness (unique genera per library) and uniqueness (taxa exclusive to each library within the dataset). Sequencing depth was categorized (<5 Gb = “low”, 5-20 Gb = “medium”, 20-100 Gb = “high”, >100 Gb = “very high”), cell size was categorized (<100 micron = “small”, 100-300 micron = “medium”, >300 micron = “large”), lifestyle (mixotrophy, heterotrophy), other features (“shell”, “microanaerobic”, “shell and algae symbionts”, “other”) and used to normalize both richness and uniqueness. We then fit separate Ordinary Least Squares models for normalized richness and uniqueness, incorporating the factors host taxonomy, cell size, other_features, sampling site, sequencing depth, and lifestyle as categorical predictors. Type II ANOVA was applied to each model, and partial eta-squared values were used to estimate each predictor’s effect size. This approach allowed us to rank factors by their influence on microbial diversity metrics.

### Screening for viruses and viral host prediction

To identify any viral sequences within the protist single cell sequence data, geNomad (v1.6.0)^61^ was used with default settings. Contigs that were predicted as viral were then subject to completeness and contamination estimate with CheckV (v1.02)^62^. Host prediction was performed with iPHoP (v1.3.3)^45^ on contigs that were predicted as viral genomes with high completeness and no contamination. To identify giant virus metagenome assembled genomes (GVMAGs) and unbinned contigs that had a length of at least 50kb were subject to classification with gvclass (v0.9) (https://github.com/NeLLi-team/gvclass)^34^. In brief, 9 conserved giant virus orthologous groups (GVOGs)^63^ were identified using hmmsearch, extracted, and used as query for a diamond blastp search (v2.1.3)^64^ against a database of the respective GVOG built from a representative set of bacteria, archaea, eukaryotes and viruses. The top 100 blastp hits were extracted, combined with the query sequence, aligned with mafft (v7.490; -linsi)^65^, trimmed with trimal (v1.4; -gt 0.1)^66^ and used to build a phylogenetic tree with IQtree(v2.3.0; LG4X)^67^. The nearest neighbor in the tree was identified using branch length and the existing taxonomic string for that reference genome was then taken into account for the final classification result. For a successful classification, the taxonomic strings from all identified nearest neighbors were compared at the different taxonomic levels (genus, family, order, class, phylum) to yield the final classification at the lowest taxonomic level on which all nearest neighbors were in agreement.

### Eukaryotic phylogenomics

To reconstruct a phylogenetic tree showing the position of our focal taxa within the eukaryotic tree of life, we used our phylogenomic pipeline PhyloToL^68^. Gene trees for 391 gene families that are highly conserved across eukaryotes and present in at least four out of five major eukaryotic clades were produced. We chose 250 taxa from all major eukaryotic clades as well as bacteria and archaea, with an even distribution of 25 taxa per major clade. In addition, one representative of each of our focal species was added. PhyloToL produces multi-sequence alignments using Guidance v2.0^69^ and builds gene trees using RAxML (PROTGAMMALG)^70^. In addition, we generated a supermatrix using the alignment concatenation option of PhyloToL and inferred a species tree with RAxML (PROTGAMMALG).

### Bacterial and viral phylogenomics

An initial bacterial species tree was built with New Simple Genome Tree (nsgtree v.0.4.0, https://github.com/NeLLi-team/nsgtree) from all MAGs that had at least 5/56 single copy marker genes of the UNI56 set of markers^58^ together with a representative set of genomes from the GTDB database^63^. GVOGs were identified with hmmsearch (v.3.3.2), aligned with mafft(v.7.508)^65^ and trimmed with trimal (v1.4)^66^. Clades of known symbionts were then selected and extracted from the tree, additional genomes (1 per family) were added to the selected clades and separate species trees were built.

To infer a species tree for the Nucleocytoviricota, GVMAGs and a reference dataset of published giant virus genomes^63^ were combined and nsgtree was employed with the set of phylogenetic markers GV0G8 (GVOGm0013, GVOGm0022, GVOGm0023, GVOGm0054, GVOGm0172, GVOGm0461, GVOGm0760, GVOGm0890)^63^; genomes that had less than 4 out of 8 GVOGs were removed. GVOGs were identified with hmmsearch (v.3.3.2), aligned with mafft(v.7.508)^65^ and trimmed with trimal (v1.4)^66^. Only GVMAGs and reference giant virus genomes with at least four of the seven GVOGs and with no more than four copies of any of the GVOG8 were selected, totalling 82 GVMAGs with assembly sizes ranging from 28kb to 1.43Mb. The final tree was calculated with IQtree (v.2.1.11)^67^ and visualized with iTOL (v.6)^71^.

## Acknowledgements

The work conducted by the U.S. Department of Energy Joint Genome Institute (https://ror.org/04xm1d337), a DOE Office of Science User Facility, is supported by the Office of Science of the U.S. Department of Energy operated under Contract No. DE-AC02-05CH11231 and through NSF grants DEB-2230391 and OCE-1924570, and NIH award R15HG010409 to LAK.

---

**Supplementary table 1. Overview of the sampled protists and sequencing strategy.**

**Supplementary table 2. Assembly statistics of metagenome assembled genomes recovered in this study.**

**Supplementary table 3. Detailed characterization of giant virus metagenome assembled genomes recovered in this study.**

**Supplementary table 4. Assembly statistics of Virophages identified in this study.**

**Supplementary table 5. Other DNA viruses discovered in this study.**

**Supplementary table 6. Results from crossblock filtering**

